# Metabolic proof-reading in *Plasmodium berghei*: essentiality of phosphoglycolate phosphatase

**DOI:** 10.1101/495473

**Authors:** Lakshmeesha Kempaiah Nagappa, Pardhasaradhi Satha, Thimmaiah Govindaraju, Hemalatha Balaram

**Affiliations:** From the Molecular Biology and Genetics Unit and Jawaharlal Nehru Centre for Advanced Scientific Research (JNCASR), Bengaluru, Karnataka, INDIA; From the Bioorganic Chemistry Laboratory, New Chemistry Unit, Jawaharlal Nehru Centre for Advanced Scientific Research (JNCASR), Bengaluru, Karnataka, INDIA

**Author notes:** To whom correspondence should be addressed: Hemalatha Balaram, Molecular Biology and Genetics Unit, Jawaharlal Nehru Centre for Advanced Scientific Research (JNCASR), Jakkur P.O., Bengaluru, Karnataka, 560064, India.

**Keywords:** *P berghei*, phosphoglycolate phosphatase, 2-phosphoglycolate, 2-phospho L-lactate, 4-phosphoerythronate, metabolic proof-reading, detoxification

## Abstract

***Plasmodium falciparum* (Pf):** 4-nitrophenylphosphatase was earlier shown to be involved in vitamin B1 metabolism by Knöckel *et al.*, (Mol. Biochem. Parasitol. 2008, **157**, 241-243). An independent BLASTp search showed that the protein had significant homology with phosphoglycolate phosphatase from mouse, human and yeast, and prompted us to re-investigate the biochemical properties of the recombinant *Plasmodium* enzyme. Owing to the insoluble nature of the Pf enzyme, an extended substrate screen and biochemical characterization was performed on its *P. berghei* (Pb) homolog that led to the identification of 2-phosphoglycolate and 2-phospho L-lactate as the relevant physiological substrates. 2-phosphoglycolate is known to be generated during repair of damaged DNA ends whereas, 2-phospho L-lactate is a product of pyruvate kinase side reaction. These metabolites are potent inhibitors of the key glycolytic enzymes, triosephosphate isomerase and phosphofructokinase, and hence clearance of these toxic metabolites is vital for cell survival and functioning. Gene knockout studies conducted in *P. berghei* revealed the essential nature of this conserved ‘metabolic proof-reading enzyme’.

---

Haloacid dehalogenase superfamily (HADSF) is a large family of enzymes consisting mainly of phosphatases and phosphotransferases, that are both intracellular and extracellular in nature. These enzymes are characterized by the presence of a core Rossmanoid-fold and a cap-domain (1, 2). Studies on HADSF members have focused on identifying their physiological substrates by screening a wide range of metabolites that include sugar phosphates, lipid phosphates, nucleotides as well as phosphorylated amino acids and co-factors. This approach has helped understand the physiological relevance of these enzymes in various cellular processes such as cell wall synthesis, catabolic and anabolic pathways, salvage pathways, signaling pathways and detoxification (3–13). Apart from dephosphorylating metabolites, HADSF members have also been know to dephosphorylate proteins and such members are characterized by the absence of the cap domain (1, 2). A large scale study reported by Huang *et al*., has identified a HADSF member from *Salmonella enterica* that catalyzes dephosphorylation of more than 100 phosphorylated substrates (5). This extended substrate specificity is a common observation in HADSF members and often leads to a confounding situation where determining the physiological substrate of such promiscuous enzymes becomes a challenging task.

Recent studies have identified and characterised HADSF members from the apicomplexan parasite, *Plasmodium* (4, 10, 13–16). HADSF members from *Plasmodium* have been found to be involved in processes that lead to the development of resistance to the drug fosmidomycin, which inhibits isoprenoid biosynthesis (4). Also, these enzymes show considerable activity towards nucleotide monophosphates and phosphorylated co-factors, and generic substrates such as p-nitrophenylphosphate (pNPP) and *β*-glycerophosphate. A HADSF member that was annotated as 4-nitrophenylphosphatase from *P. falciparum* (gene id. PF3D7_0715000) was characterized by Knöckel *et al.*, (2008) and was proposed to be involved in dephosphorylation of thiamine monophosphate, the precursor of the active form of vitamin B1 (thiamine pyrophosphate). *In vitro* assays on the purified recombinant enzyme showed that this protein displayed similar specific activities towards thiamine monophosphate and other substrates (ADP, ATP, CTP, G-6-P, F-6-P and PLP) (15). An independent BLASTp search conducted by us revealed that this protein sequence has significant homology (28-30 %) with phosphoglycolate phosphatase (PGP) from yeast, human and mouse (Fig. 1). The (His)_6_-tagged recombinant *P. falciparum* (Pf) 4-nitrophenylphosphatase when expressed in *Escherichia coli*, was found to be completely insoluble. However, *P. berghei* (Pb) 4-nitrophenylphosphatase (gene id. PBANKA_1421300) (referred to as PbPGP here-onwards) that shares 69.6 % identity (Fig. 1B) with its *Pf* homolog, expressed in the soluble form in *E. coli* and could be purified to homogeneity. Here, we report on the biochemical characterization and essentiality of PbPGP. An extended substrate screen identified 2-phosphoglycolate and 2- phospho L-lactate as relevant physiological substrates in addition to the generic substrates pNPP and *β*-glycerophosphate. Attempts at gene ablation showed that PbPGP gene cannot be disrupted in *P. berghei*, despite the loci being non-refractory for genetic recombination. Our studies on PbPGP establish the essential physiological nature and biochemical function of this conserved cytosolic enzyme, and suggest that drugs that specifically inhibit parasite phosphoglycolate phosphatase can be promising anti-malarial agents.

**Figure 1:**
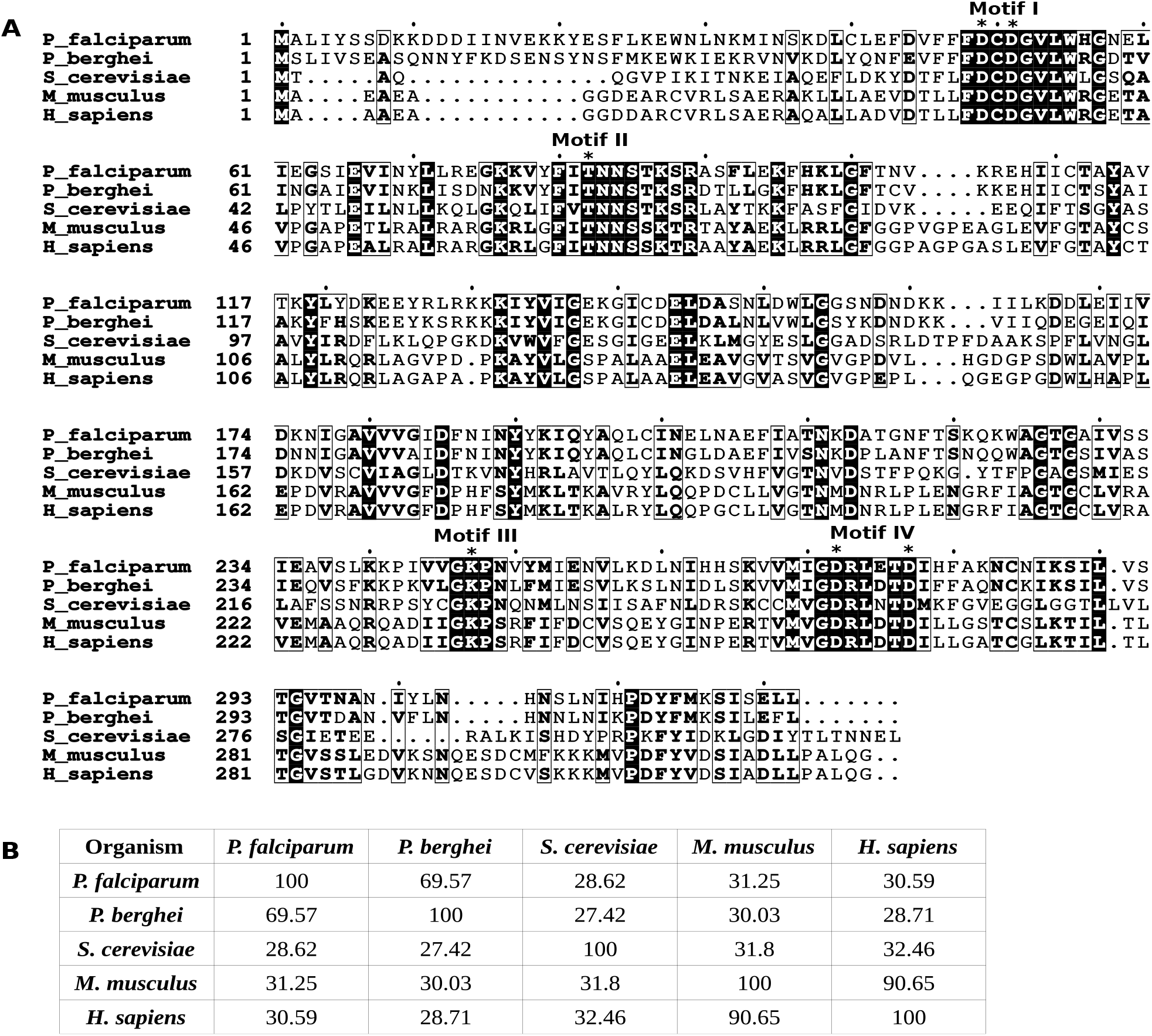
Multiple sequence alignment of phosphoglycolate phosphatase protein sequences. (A) Clustal omega alignment of phosphoglycolate phosphatase from *Plasmodium falciparum* (P_falciparum), *Plasmodium berghei* (P_berghei), *Saccharomyces cerevisiae* (S_*cerevisiae*), *Mus musculus* (M_musculus) and human (H_sapiens). Residues of the conserved HAD motifs involved in catalysis are indicated by *. (B) Percentage identity matrix showing the extent of homology between the sequences.

## RESULTS

### Biochemical characterization of recombinant PbPGP

BLASTp analysis of Pf and Pb PGP protein sequences showed 28-30 % sequence homology with phosphoglycolate phospatase, a conserved protein, present across eukaryotes from yeast to mouse including humans, involved in metabolic proof-reading (Fig. 1A). The Pf and Pb protein sequences show 69.6 % identity (Fig. 1B).

Upon expression of C-terminal (His)_6_-tagged PfPGP in Rosetta DE3 pLysS strain of *E. coli*, the protein was found to be present completely in the insoluble fraction (Fig. S1A). This was unlike the strep-tagged PfPGP that was reported to be present in small quantities in the soluble fraction and hence amenable to purification. Therefore, we made use of the protein solubility prediction software PROSOII and found that homologues of PfPGP from other Plasmodia were predicted to be soluble (Fig. S1B). Hence, the previously uncharacterized *P. berghei* homolog was chosen for further biochemical studies and physiological investigations. PbPGP was expressed in the *E. coli* strain Rosetta DE3 pLysS and purified to homogeneity by Ni-NTA affinity chromatography (Fig. S1C) followed by size-exclusion chromatography (Fig. 2A).

**Figure 2:**
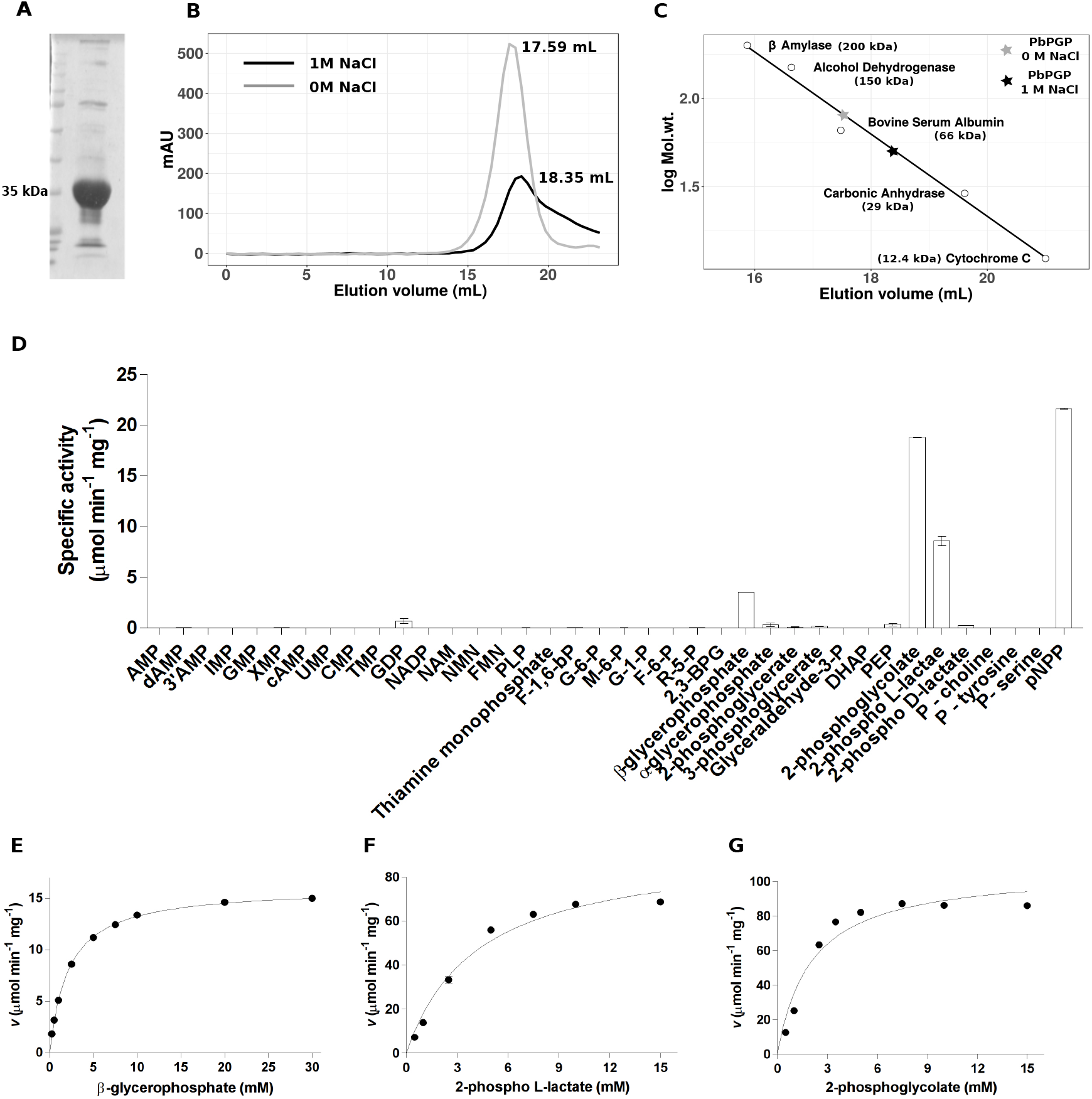
Purification and biochemical characterization of PbPGP. (A) SDS-PAGE of PbPGP purified using Ni-NTA affinity followed by size-exclusion chromatography. (B and C) Determination of oligomeric state of PbPGP. (B) Elution profile of PbPGP in the presence and absence of 1 M NaCl and (C) molecular mass calibration curve with elution volumes of PbPGP in the absence and presence of NaCl interpolated. (D) Screen for potential substrates of PbPGP. Mean specific activity values are provided for each substrate and error bars represent SD (n=2). AMP, adenosine 5’ monophosphate; dAMP, deoxy adenosine 5’ monophosphate; 3’AMP, adenosine 3’ monophosphate; IMP, inosine 5’ monophosphate; GMP, guanosine 5’ monophosphate; XMP, xanthosine 5’ monophosphate; cAMP, 3’ 5’ cyclic AMP; UMP, uridine 5’ monophosphate; CMP, cytidine 5’ monophosphate; TMP, thymidine 5’ monophosphate; GDP, guanosine diphosphate; NADP, nicotinamide adenine dinucleotide phosphate; NAM, nicotinic acid mononucleotide; NMN, nicotinamide mononucleotide; FMN, flavin mononucleotide; F-1,6-bp, fructose 1,6-bisphosphate; G-6-P, glucose 6-phosphate; M-6-P, mannose 6-phosphate; G-1-P, glucose 1-phosphate; F-6-P, fructose 6-phosphate; R-5-P, ribose 5-phosphate; 2,3-BPG, 2,3-bisphosphoglycerate; DHAP, dihydroxyacetone phosphate; PEP, phosphoenolpyruvate. (E-G) Substrate concentration vs. specific activity plots fit to Michaelis-Menten equation for *β*-glycerophosphate, 2-phospho L-lactate and 2-phosphoglycolate. Substrate titration experiment was conducted in two technical replicates containing two biological replicates each. Plots from one technical replicate are shown. Each data point represents mean specific activity value and error bars represent SD (n=2).

PbPGP on analytical gel filtration using Sephacryl S-200 coulmn showed a mass of about 78 kDa, whereas the theoretical mass is 37 kDa, indicating that the protein is a dimer (Fig. 2B and C). When further analyzed in the presence of 1 M NaCl, there was a shift in oligomeric state of the protein from dimer, towards monomer suggesting that the oligomers are held by electrostatic interactions (Fig. 2B and C).

A total of 38 compounds were screened as possible substrates for PbPGP. Although the enzyme displayed very low activity towards nucleotides and sugar phosphates as reported for PfPGP by Knöckel *et al.*, (15) a novel observation was made as a consequence of our extended substrate screen. PbPGP showed very high activity on 2-phosphoglycolate and 2-phospho L-lactate in addition to the generic substrates pNPP and *β*-glycerophosphate (Fig. 2D). It should be noted that the enzyme was stereospecific for 2-phospho L-lactate and showed no activity on 2-phospho D-lactate.

### Kinetic studies on PbPGP

PbPGP showed maximum activity at pH 7.0 and preferred Mg^2+^ as co-factor over other divalent cations (Fig. S2A and B). The substrate saturation plots for *β*-glycerophosphate, 2-phosphoglycolate and 2-phospho L-lactate were hyperbolic and were fit to Michaelis-Menten equation to obtain the kinetic parameters such as *K*_m_ and *V*_max_ (Fig. 2E-G and Table 1). PbPGP has higher *K*_m_ value for 2-phosphoglycolate (3.3 and 11.4 fold) and 2-phospho L-lactate (27.4 and 6.4 fold) when compared with that of murine PGP and yeast Pho13. The *k*_cat_ value for PbPGP for 2-phosphoglycolate is 11.4 and 3.9 fold higher and for 2-phospho L-lactate is 37 and 8.9 fold higher when compared to that of murine and yeast homologs, respectively. The catalytic efficiency (*k*_cat_/*K*_m_) for 2-phosphoglycolate was 3.5 fold higher and 2.9 fold lower when compared with its murine and yeast homologs respectively. With 2-phospho L-lactate as substrate, the parasite enzyme has similar catalytic efficiency as its murine and yeast homologs.

**Table 1:** Kinetic parameters of *P. berghei* PGP compared with that of homologs from yeast and mouse.

| Substrate | Kinetic parameters of PbPGP |  |  |  |
| --- | --- | --- | --- | --- |
| | $K_m$ ($\mu\text{M}$) | $V_{\max}$ ($\mu\text{mol min}^{-1} \text{mg}^{-1}$) | $k_{\text{cat}}$ ($\text{s}^{-1}$) | $k_{\text{cat}}/K_m$ ($\text{M}^{-1} \text{s}^{-1}$) |
| $\beta$ - glycerophosphate | $2110 \pm 11$ | $16.2 \pm 0.14$ | $10.18 \pm 0.08$ | 4827 |
| 2 - phosphoglycolate | $2526 \pm 494$ | $119.5 \pm 12.5$ | $75.15 \pm 7.86$ | 29747 |
| 2 - phospho L - lactate | $4773 \pm 574$ | $107 \pm 13.2$ | $67.34 \pm 8.31$ | 14108 |
| Kinetic parameters of Murine PGP<sup>#</sup> |  |  |  |  |
| 2 - phosphoglycolate | $766 \pm 68$ | $11.34^*$ | $6.56 \pm 0.44$ | 8564 |
| 2 - phospho L - lactate | $174 \pm 55$ | $3.14^*$ | $1.82 \pm 0.34$ | 10480 |
| Kinetic parameters of *S. cerevisiae* Pho13<sup>#</sup> |  |  |  |  |
| 2 - phosphoglycolate | $221 \pm 13$ | $32.87^*$ | $19.0 \pm 0.44$ | 85700 |
| 2 - phospho L - lactate | $747 \pm 135$ | $13.09^*$ | $7.57 \pm 1.04$ | 10113 |

Data represents mean ± S.E.M (N=2)
<sup>#</sup>Values taken from Collard et al., 2016
\* $V_{\max}$ was calculated using $k_{\text{cat}}$ values from Collard et al., 2016. Molecular mass values of 34540.68 Da and 34624.58 Da for murine PGP and Pho13, respectively was used in the calculation.

### Probing the essentiality of PbPGP and localization in P. berghei

pJAZZ linear knockout vector for PbPGP was generated by following the strategy described by Pfander *et al.*, (17). Drug resistant parasites were not obtained in the first transfection attempt. In the second attempt, though drug resistant parasites were obtained, genotyping by PCR revealed non-specific integration of the marker cassette. These parasites were positive by PCR for both the PbPGP gene and the hDHFR marker but were negative for specific 5’ and 3’ integration PCRs (Fig. S3). Since it was not possible to obtain knockout parasites, a conditional knockdown (at the protein level) strategy was employed by tagging the gene for PbPGP with a regulatable fluorescent affinity tag (RFA) where stability of the fusion protein is conditional to the binding of the small molecule trimethoprim. The conditional knockdown vector was also generated by following the recombineering strategy and validated by PCR (Fig. 3A-F). Transgenic parasites were obtained in the first transfection attempt itself and genotyping by PCR showed the presence of a single homogenous population with correct insertion of RFA tag (Fig. 3F). Nevertheless, it was observed that the reduction in the levels of RFA-tagged protein upon removal of TMP, varied between 30-60 % across experiments and complete knockdown could not be achieved (Fig. 3G-I).

**Figure 3:**
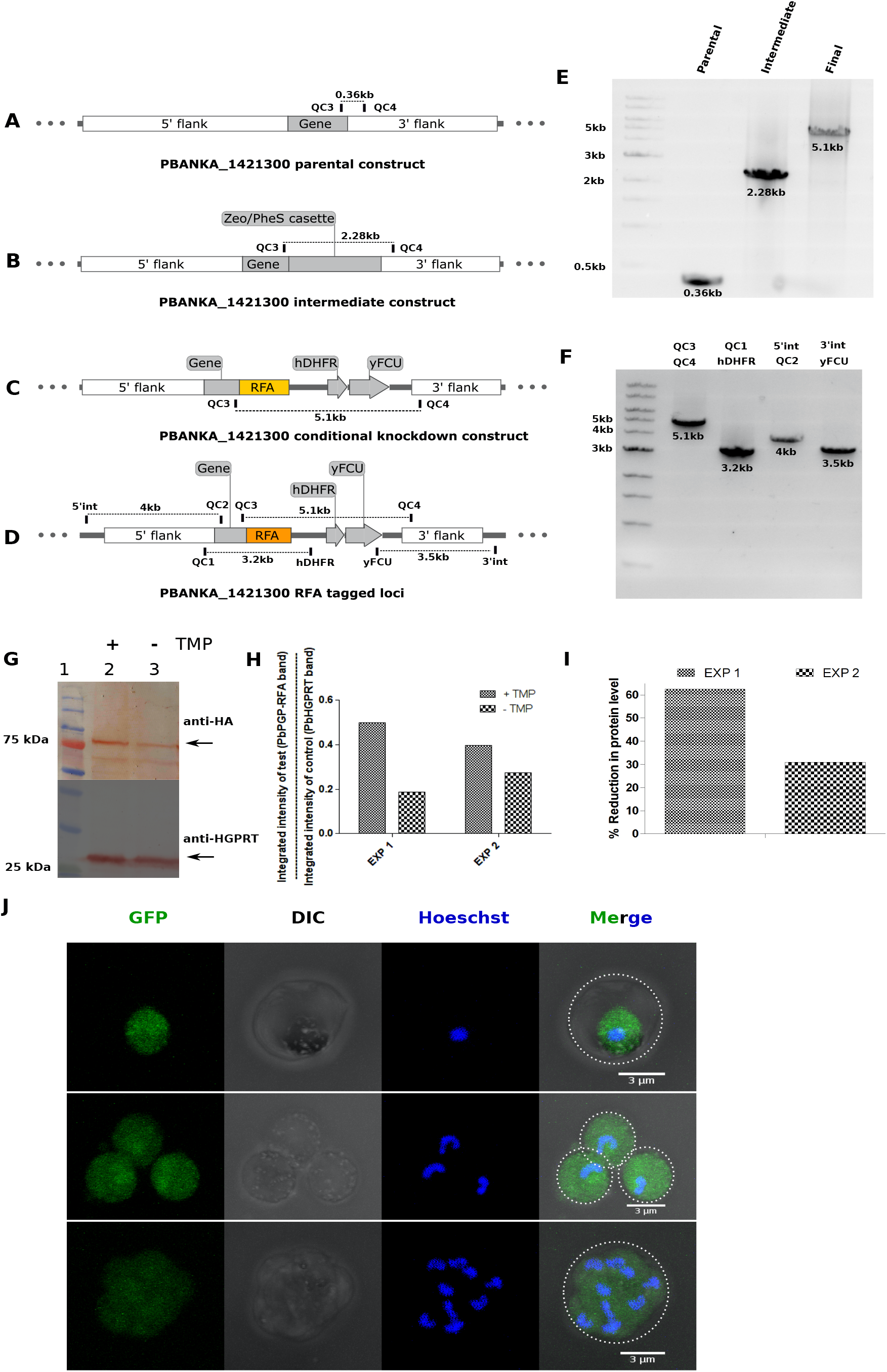
Conditional knockdown of PbPGP and localization in *P. berghei*. (A-D) Schematic representation of PbPGP parental, intermediate and final RFA tagging constructs and PbPGP loci after integration. Oligonucleotide primers are indicated by vertical bars and expected PCR-product size is represented by line between specific primer pairs. (E) PCR confirmation of parental, intermediate and final RFA-tagging construct. (F) Genotyping of the strain for integration of cassette in the correct loci. Primer pairs used are mentioned on top of the panel. (G) Western blot analysis of cell lysates of RFA-tagged parasites from mice fed with/without trimethoprim (TMP) (30 mg in 100 mL) for 6 days. The experiment was performed twice (Exp1 and Exp2) and blot from one experimental replicate is shown. Top panel is the blot probed with anti-HA antibody and the arrow mark indicates the RFA tagged PbPGP protein. The bottom panel was probed with anti-PfHGPRT antibody. (H) Ratio of the intensity of RFA tagged PbPGP to the intensity of control HGPRT. (I) Reduction in PbPGP levels upon removal of TMP relative to the levels in the presence of TMP. Protein levels under ‘+ TMP’ condition was taken as 100 %. (J) Localization of PbPGP using RFA-tagged parasites grown in ‘+ TMP’ condition. The erythrocyte boundary is indicated by white dotted line in the merge panel.

The transgenic RFA-tagged *P. berghei* parasites were employed to determine localization of PbPGP and upon microscopic observation a cytosolic GFP signal was observed in all the intra-erythrocytic stages (Fig. 3J).

## DISCUSSION

Earlier Knöckel *et al.*, had performed a TBLASTN search and identified a potential 4-nitrophenylphosphatase in *P. falciparum*. The authors had proposed a novel role for this HADSF member and suggested involvement in vitamin B1 homeostasis (15). We found the *P. falciparum* 4-nitrophenylphosphatase sequence to have homology with human, mouse and yeast phosphoglycolate phosphatases. An extended substrate specificity screen of the recombinant *P. berghei* enzyme revealed that, indeed this protein is phosphoglycolate phosphatase, which is mainly involved in detoxification, having very high activity on 2-phosphoglycolate and 2-phospho L-lactate with no activity on thiamine monophosphate. 2- phosphoglycolate is reported to be formed during repair of free radical mediated damage of DNA ends (18) and accumulation of this metabolite in the cell, leads to inhibition of the key glycolytic enzyme triosephosphate isomerase (TIM) (Fig. 4). Studies on phosphoglycolic acid phosphatases from yeast and mouse have demonstrated that this enzyme also performs metabolic proof-reading by catabolizing the substrates 2-phospho L-lactate and 4-phosphoerythronate which are products of enzymatic side reactions. Activity of PbPGP on 4-phosphoerythronate could not be tested due to non availability of the compound. 2-phospho L-lactate, generated by phosphorylation of L-lactate by pyruvate kinase, is known to inhibit phosphofructokinase and 4-phosphoerythronate, which is a product of GAPDH side reaction, is known to inhibit 6-phosphogluconate dehydrogenase (Fig. 4) (19). Due to the detrimental effect of these metabolites, it becomes essential to clear the cell of these metabolic toxins. This is reflected upon by the fact that phosphoglycolate phosphatase is an essential gene in mouse (20). Also, in *Arabidopsis*, knockout of PGLP1 isoform leads to impaired postgermination development of primary leaves (21).

**Figure 4:**
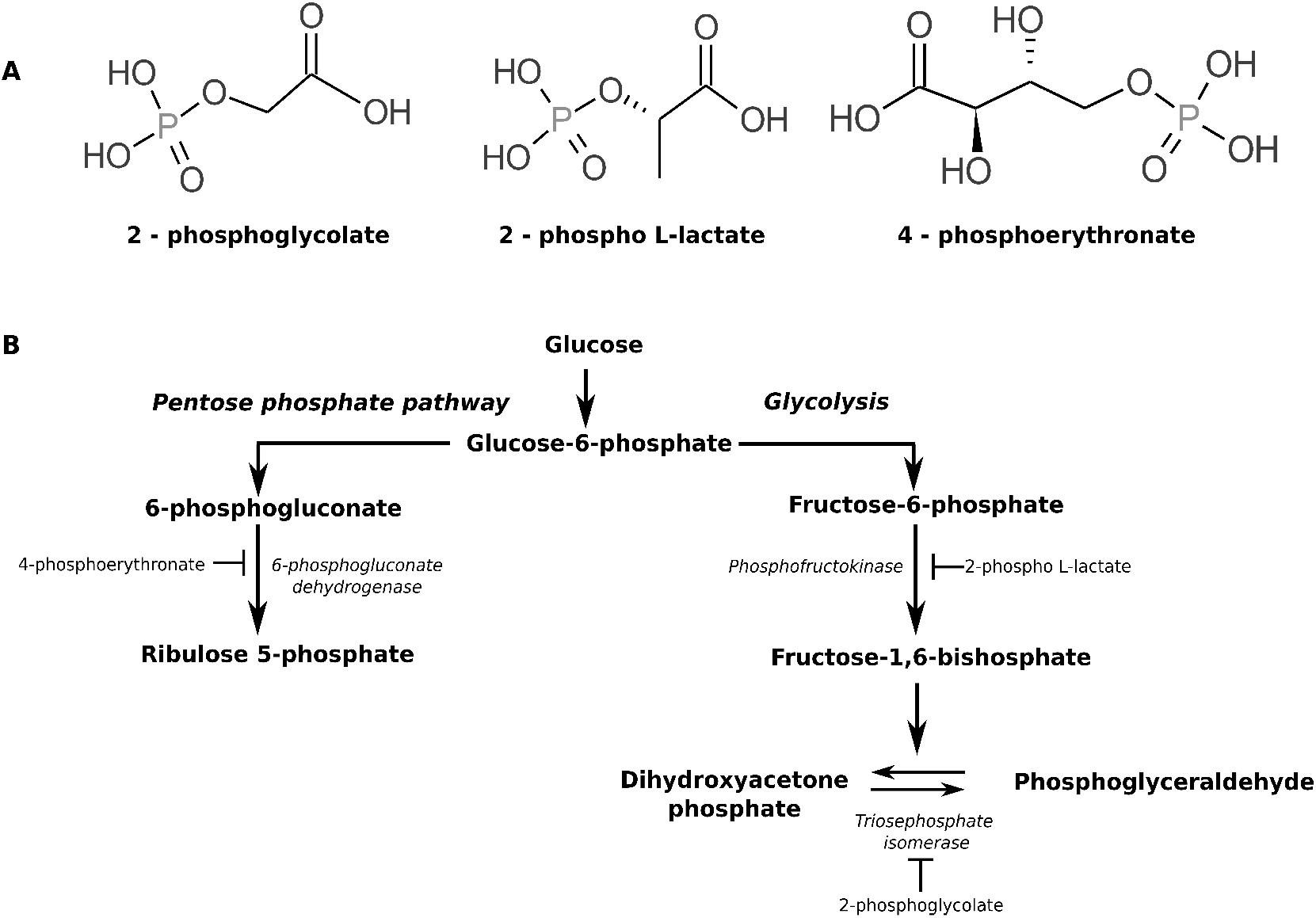
Metabolites that are substrates for phosphoglycolate phosphatase (A) and their inhibition of key metabolic pathways (B).

*Plasmodium* in its intra-erythrocytic stages experiences very high levels oxidative stress (22) leading to increased ROS production that can damage its DNA, the repair of which will result in generation and accumulation of 2-phosphoglycolate. The parasite undergoes lactic acid fermentation and is known to secrete large amounts of lactate into the medium, most of which is L-lactate (93-94 %), in addition to a small proportion of D-lactate (6-7 %) that is known to be produced through the methylglyoxal pathway (23). This lactate can accumulate and be phosphorylated in the cell to give rise to 2-phospholactate. Surprisingly, Dumont *et al.*, show that upon addition of D-lactate to Δ*PfPGP* cells, intracellular levels of 2-phospholactate spike up and growth is retarded. The authors speculate that although L-lactate is the major isomer that is produced in the parasite, it is D-lactate that is phosphorylated (16) and accumulates in Δ*PfPGP* cells. Our results on the purified enzyme clearly show that PbPGP acts only on 2-phospho L-lactate and not on 2-phospho D-lactate (Fig. 2D). We further validated this by performing enzyme assays on 1 mM 2-phospho L-lactate in the presence or absence of 2 mM 2-phospho D-lactate. There was no significant change in specific activity clearly showing that 2-phospho D-lactate does not bind to the enzyme (Fig. S2C). This observation is consistent with that of the murine homolog of PbPGP that also acts only on 2-phospho L-lactate (19). It has to be noted that recombinant human pyruvate kinase M2 isoform has been shown to phosphorylate L-lactate leading to the production of 2-phospho L-lactate. Also, 2-phospho L-lactate was shown to inhibit phosphofructokinase-2 activity in crude lysates of HCT116 cells and activity of recombinant phosphofructokinase-fructose 1,6 bisphosphatase (PFKFB) isozymes, PFKFB3 and PFKFB4 (19).

In *Plasmodium*, where glycolysis is the sole source of ATP in asexual stages (24), the parasite cannot afford inhibition of its critical enzymes such as PFK and TIM due to accumulation of toxic metabolites. Hence, having a metabolic proofreading/detoxifying enzyme becomes vital for its survival. Inability to obtain knockout parasites indicates essentiality of this protein for parasite survival during asexual stages. To rule out the possibility of the loci being refractory for genetic recombination regulatable fluorescent affinity (RFA) tagging was attempted and a homogenous poplulation of transfectants with RFA-tag integrated at the right loci was obtained. Having established that the loci is amenable for genetic manipulation, conditional knockdown strategy at protein level making use of the RFA tag (25) was adopted. Conditional knockdown at the protein level showed only 30-60 % reduction and hence parasites were viable. Similar observation has been described for yoelipain where the authors were neither able to knockout nor achieve significant knockdown of protein levels to see growth difference. Therefore, it was concluded that the gene was essential during intra-erythrocytic stages (26). Our results are similar and indicate essentiality of PbPGP in asexual stages. This conclusion on the gene essentiality of PbPGP is in agreement with the findings of Dumont *et al.*, on PfPGP where, Δ*pfpgp* parasites show growth defect (16). Further biochemical and structural studies on PGP could pave way for rational design of inhibitors with potent anti-malarial activity.

## MATERIALS

All chemicals, molecular biology reagents, and media components were from Sigma Aldrich, New England Biolabs, Gibco, Invitrogen and US biochemicals, USA; SRL, Spectrochem and Hi-media, India. *E. coli* strain XL-1 blue, expression strain Rosetta (DE3) pLysS, and plasmids pET22b, and pET23d were from Novagen. The pJAZZ library clone (PbG02_B-53b06), plasmids pSC101BAD, R6K Zeo/pheS, and GW_R6K_GFP-mut3, and *E. coli* pir strains were procured from PlasmoGem, Sanger Institute, UK. *E. coli* TSA cells were from Lucigen. Plasmid pGDB was a kind gift from Dr. Vasant Muralidharan, University of Georgia, USA. The *P. falciparum* 3D7 strain and *P. berghei* ANKA strain were procured from MR4. Amaxa 4D nucleofector and P5 Nucleofection kit were from Lonza, Germany. Gene sequencing of the various plasmid constructs was by Sanger sequencing method. Sequences of oligonucleotides used are provided in the section ‘Supplementary information’ (Table S1).

## EXPERIMENTAL PROCEDURES

### Bio-informatic analysis

PfPGP (Plas-moDB gene ID PF3D7_0715000) protein sequence obtained from PlasmoDB database was subjected to homology search against the non-redundant database at NCBI using the BLASTp algorithm. Clustal Omega (27) was used to generate multiple sequence alignment. ProsoII (28) was employed to predict solubility of proteins upon heterologous expression in *E. coli* system.

### Cloning expression and purification of Pf-PGP and PbPGP

PfPGP and PbPGP were cloned in pET23d and pET22b respectively and expressed in Rosetta DE3 pLysS strain of *E. coli.* The methods are described in detail in ‘Supplementary information’.

### Determination of oligomeric state

Oligomeric state of PbPGP was determined by analytical size-exclusion chromatography using Sephacryl S-200 (1 cm × 30 cm) column attached to an AKTA Basic HPLC system. The column was equilibrated using 100 mM Tris HCl, pH 7.4 and 100 mM KCl at 0.8 mL min^-1^ flow rate and calibrated using the molecular weight standards; *β*-amylase (200 kDa), alcohol dehydrogenase (150 kDa), bovine serum albumin (66 kDa), carbonic anhydrase (29 kDa) and cytochrome C (12.4 kDa). 100 *μ*L of PbPGP at 1 mg mL^-1^ concentration was injected into the column and eluted with equilibration buffer with monitoring at 280 nm. The molecular mass of PbPGP was estimated by interpolating the elution volume on a plot of logarithm of molecular weight standards on the Y-axis and elution volume on the X-axis. Gel-filtration was performed with and without NaCl in the equilibration buffer.

### Synthesis of 2-phospholactate

Synthesis of both D and L phospholactate was carried out following available procedure (19). The details of the protocol and characterization of the molecules are provided in ‘Supplementary information’.

### Enzyme assays

A comprehensive substrate screen comprising of various classes of molecules such as, nucleoside phosphates, sugar phosphates, co-enzymes, amino acid phosphates, etc. was performed. The assay was carried out in 100 mM Tris HCl, pH 7.4, 2 mM substrate, 1 mM MgCl_2_ in a volume of 100 *μ*L. The reaction mix was pre-incubated at 37 °C for 1 minute, the assay was initiated using 2 *μ*g enzyme and the reaction was allowed to proceed at 37 °C for 5 minutes. The reaction was stopped by the addition of 20 *μ*L of 70 % trichloroacetic acid (TCA) and 1 mL of freshly prepared Chen’s reagent (water, 6 N sulphuric acid, 2.5 % ammonium molybdate and 10 % L-ascorbic acid mixed in the ratio 2:1:1:1) was added, mixed thoroughly and incubated at 37 °C for 1.5 hours. The color developed was measured against blank (reaction mix to which enzyme was added after addition of TCA) at 820 nm. Specific activity was calculated using the *ϵ* value of 25000 M^-1^ cm^-1^.

pH optimum of PbPGP was determined by performing the assay in a mixed buffer containing 50 mM each of glycine, MES, Tris at different pH, 1 mM MgCl_2_ and 1 mM pNPP as substrate in 100 *μ*L volume. The reaction mix was pre-incubated at 37 °C for 1 minute and the assay was initiated using 0. 2 *μ*g enzyme and the reaction was allowed to proceed at 37 °C for 2 minutes, stopped using TCA and processed using Chen’s reagent as described above.

Preferred divalent metal ion was identified by using 10 mM pNPP as substrate and different salts such as MgCl_2_, MnCl_2_, CaCl_2_, CuCl_2_, and CoCl_2_ at a final concentration of 1 mM in a 250 *μ*L reaction mix containing 50 mM Tris HCl, pH8. The reaction was initiated with 0.26 *μ*g of enzyme and conversion of pNPP to p-nitrophenol was continuously monitored at 405 nm at 37 °C temperature. Slope of the initial 20 seconds of the progress curve was used to calculate specific activity using an *ϵ* value of 18000 M^-1^ cm^-1^.

### Kinetic studies

*K*_m_ values for 2-phosphoglycolate, 2-phospho L-lactate and *β*-glycerophosphate was determined by measuring initial velocity at varying substrate concentrations ranging from 0.5 mM to 15 mM for 2-phosphoglycolate and 2-phospho L-lactate and 0.25 mM to 30 mM for *β*-glycerophosphate. The concentration of MgCl_2_ was fixed at 5 mM with the reaction buffer being 200 mM Tricine-NaOH, pH 7.4. The reaction in a volume of 100 *μ*L was initiated with 1.89 *μ*g of enzyme, allowed to proceed at 37 °C for 2 minutes, stopped using TCA and processed using Chen’s reagent as described above. Specific activity was plotted as a function of substrate concentration and the data points were fitted to the Michaelis-Menten equation using GraphPad prism V5 to determine the kinetic parameters (29).

### Generation of P. berghei transfection vectors

The library clone for *P. b*erghei PGP (PbG02_B-53b06) was obtained from PlasmoGem. The procedure for knockout and tagging construct generation was as described earlier (17, 30) and is provided in detail in ‘Supplementary information’.

### Cultivation and transfection of P. berghei

Male/female BALB/c mice aged 6-8 weeks were used for cultivation and transfection of *P. berghei.* Glycerol stock of wild type *P. berghei ANKA* parasites was injected into a healthy male BALB/c mouse. The parasitemia was monitored by microscopic observation of Giemsa stained smears of blood drawn from tail snip. Transfection of the parasites was done by following the protocol described by Janse *et al.*, (31), using Amaxa 4D nucleofector (P5 solution and FP167 programme) followed by injection into 2 mice. For PbPGP knockout, drug resistant parasites were selected for by feeding infected mice with pyrimethamine in drinking water (7 mg in 100 mL), whereas parasites with PbPGP RFA-tag were selected for by feeding infected mice with trimethoprim in drinking water (30 mg in 100 mL). Drug resistant parasites were harvested in heparin solution (200 units mL^-1^) made in RPMI-1640. Glycerol stocks were made by mixing 300 *μ*L of 30 % glycerol and 200 *μ*L of the harvested blood and stored in liquid nitrogen. Validation of the drug resistant parasites was done by PCR.

### Conditional knockdown of PbPGP in P. berghei

Glycerol stock of PbPGP RFA-tagged parasites was injected into a healthy BALB/c mouse. The parasitemia was monitored by microscopic observation of Giemsa stained smears of blood drawn from tail snip. Trimethoprim pressure was maintained throughout the growth period. Upon parasitemia reaching 5-10 %, about 500 *μ*L of infected blood was collected in 500 *μ*L of RPMI-1640 solution containing heparin. A 100 *μ*L of this parasite containing suspension was injected into a fresh mouse that was fed with trimethoprim in drinking water and a second 100 *μ*L to another mouse that was not fed with trimethoprim. Parasites were harvested from both mice after 6 days and subjected to Western blotting. The entire experiment was repeated twice.

### Localization of PGP in P. berghei

PbPGP RFA-tagged parasites were harvested in heparin solution, centrifuged at 2100 × g for 5 minutes and the supernatant discarded. The cells were resuspended in 1 × PBS containing Hoeschst 33342 (10 *μ*g mL^-1^) and incubated at room temperature for 15 minutes. Thereafter, the cells were collected, washed once with 1 × PBS, resuspended in 70 % glycerol and dispersed on poly L-lysine coated cover slips that were mounted on glass slides, sealed and stored at 4 °C. The slides were observed under oil immersion objective (100 ×) of Ziess LSM 510 Meta confocal microscope.

## Acknowledgments

This project was funded by; 1) Department of Biotechnology, Ministry of Science and Technology, Government of India. Grantnumber: BT/PR11294/BRB/10/1291/2014, BT/PR13760/COE/34/42/2015, and BT/INF/22/SP27679/2018. 2) Science and Engineering Research Board, Department of Science and Technology, Government of India. Grant number: EMR/2014/001276 and, 3) Institutional funding from Jawaharlal Nehru Centre of Advanced Scientific Research, Department of Science and Technology, India. LKN acknowledges CSIR for junior and senior research fellowships. TG acknowledges Department of Science and Technology, Government of India (Grant number: DST/SJF/CSA-02/2015-2016) for funding. PS acknowledges Science and Engineering Research Board, Department of Science and Technology, Government of India for post-doctoral fellowship (Grant number: 2017/000920). The authors thank Mr. Madhav Nayak for initial help in synthesis of phospholactate, Mrs. Suma for help in confocal microscopy and Dr. R. G. Prakash for help in animal handling.

## Conflict of interest

The authors declare that they have no conflicts of interest with the contents of this article.

## Author contributions

HB and LKN conceived the project and designed the experiments. LKN performed biochemical and physiological characterization. PS synthesized and characterized 2-phospholactate under the supervision of TG. LKN and HB wrote the manuscript.

## Ethics statement

Animal experiments involving handling of BALB/c mice were performed by adhering to standard procedures prescribed by the Committee for the Purpose of Control and Supervision of Experiments on Animals (CPCSEA), a statutory body under the Prevention of Cruelty to Animals Act of 1960 and Breeding and Experimentation Rules of 1998, Constitution of India. The current study (project no. HB006/201/CPCSEA) was approved by Institutional animal ethics committee (IAEC) that comes under the purview of CPCSEA.

## Abbreviation

HADSF: Haloacid dehalogenase superfamily
PGP: phosphoglycolate phosphatase
pNPP: para-nitrophenylphosphate
RFA: regulatable fluorescent affinity

## REFERENCES

1. Allen K. N. and Dunaway-Mariano D. (2004) Phosphoryl group transfer: evolution of a catalytic scaffold. Trends Biochem. Sci. 29, 495–503

2. Allen K. N. and Dunaway-Mariano D. (2009) Markers of fitness in a successful enzyme superfamily. Curr. Opin. Struct. Biol. 19, 658–665

3. Burroughs, A. M., Allen, K. N., Dunaway-Mariano, D., and Aravind, L. (2006) Evolutionary genomics of the had superfamily: understanding the structural adaptations and catalytic diversity in a superfamily of phosphoesterases and allied enzymes. J. Mol. Biol. 361, 1003–1034

4. Guggisberg, A. M., Park, J., Edwards, R. L., Kelly, M. L., Hodge, D. M., Tolia, N. H., and Odom, A. R. (2014) A sugar phosphatase regulates the methylerythritol phosphate (mep) pathway in malaria parasites. Nat. Commun. 5, 4467

5. Huang, H., Pandya, C., Liu, C., Al-Obaidi, N. F., Wang, M., Zheng, L., Keating, S. T., Aono, M., Love, J. D., Evans, B., et al. (2015) Panoramic view of a superfamily of phosphatases through substrate profiling. Proc. Natl. Acad. Sci. 112, E1974–E1983

6. Kuznetsova, E., Proudfoot, M., Gonzalez, C. F., Brown, G., Omelchenko, M. V., Borozan, I., Carmel, L., Wolf, Y. I., Mori, H., Savchenko, A. V., et al. (2006) Genome-wide analysis of substrate specificities of the *Escherichia coli* haloacid dehalogenase-like phosphatase family. J. Biol. Chem. 281, 36149–36161

7. Kuznetsova, E., Nocek, B., Brown, G., Makarova, K. S., Flick, R., Wolf, Y. I., Khusnutdinova, A., Evdokimova, E., Jin, K., Tan, K., et al. (2015) Functional diversity of haloacid dehalogenase superfamily phosphatases from *Saccharomyces cerevisiae* biochemical, structural, and evolutionary insights. J. Biol. Chem. 290, 18678–18698

8. Proudfoot, M., Kuznetsova, E., Brown, G., Rao, N. N., Kitagawa, M., Mori, H., Savchenko, A., and Yakunin, A. F. (2004) General enzymatic screens identify three new nucleotidases in *Escherichia coli* biochemical characterization of sure, yfbr, and yjjg. J. Biol. Chem. 279, 54687–54694

9. Roberts, A., Lee, S.-Y., McCullagh, E., Silversmith, R. E., and Wemmer, D. E. (2005) Ybiv from *Escherichia coli* k12 is a had phosphatase. Proteins: Struct. Funct. Bioinforma. 58, 790–801

10. Srinivasan, B., Nagappa, L. K., Shukla, A., and Balaram, H. (2015) Prediction of substrate specificity and preliminary kinetic characterization of the hypothetical protein pvx_123945 from Plasmodium vivax. Exp. Parasitol. 151, 56–63

11. Titz, B., Häuser, R., Engelbrecher, A., and Uetz, P. (2007) The *Escherichia coli* protein yjjg is a house-cleaning nucleotidase in vivo. FEMS Microbiol. Lett. 270, 49–57

12. Weiss B. (2007) Yjjg, a dump phosphatase, is critical for thymine utilization by *Escherichia coli* k-12. J. Bacteriol. 189, 2186–2189

13. Guggisberg, A., Frasse, P., Jezewski, A., Kafai, N., Gandhi, A., Erlinger, S., and Odom, A. J. (2018) Suppression of drug resistance reveals a genetic mechanism of metabolic plasticity in malaria parasites. MBio 9, e01193–18

14. Srinivasan B. and Balaram H. (2007) Isn1 nucleotidases and had superfamily protein fold: in silico sequence and structure analysis. In Silico Biol. 7, 187–193

15. Knöckel, J., Bergmann, B., Müller, I. B., Rathaur, S., Walter, R. D., and Wrenger, C. (2008) Filling the gap of intracellular dephosphorylation in the *Plasmodium falciparum* vitamin b1 biosynthesis. Mol. Biochem. Parasitol. 157, 241–243

16. Dumont, L., Richardson, M. B., van der Peet, P., Dixon, M. W., Williams, S. J., McConville, M. J., Tilley, L., and Cobbold, S. (2018) The metabolic repair enzyme phosphoglycolate phosphatase regulates central carbon metabolism and fosmidomycin sensitivity in *Plasmodium falciparum*. BioRxiv, 415505

17. Pfander, C., Anar, B., Schwach, F., Otto, T. D., Brochet, M., Volkmann, K., Quail, M. A., Pain, A., Rosen, B., Skarnes, W., et al. (2011) A scalable pipeline for highly effective genetic modification of a malaria parasite. Nat. Methods 8, 1078

18. Pellicer, M. T., Nunez, M. F., Aguilar, J., Badia, J., and Baldoma, L. (2003) Role of 2- phosphoglycolate phosphatase of *Escherichia coli* in metabolism of the 2-phosphoglycolate formed in DNA repair. J. Bacteriol. 185, 5815–5821

19. Collard, F., Baldin, F., Gerin, I., Bolsée, J., Noël, G., Graff, J., Veiga-da Cunha, M., Stroobant, V., Vertommen, D., Houddane, A., et al. (2016) A conserved phosphatase destroys toxic glycolytic side products in mammals and yeast. Nat. Chem. Biol. 12, 601–607

20. Segerer, G., Hadamek, K., Zundler, M., Fekete, A., Seifried, A., Mueller, M. J., Koentgen, F., Gessler, M., Jeanclos, E., and Gohla, A. (2016) An essential developmental function for murine phosphoglycolate phosphatase in safeguarding cell proliferation. Sci. Rep. 6, 35160

21. Schwarte S. and Bauwe H. (2007) Identification of the photorespiratory 2-phosphoglycolate phosphatase, pglp1, in Arabidopsis. Plant Physiol. 144, 1580–1586

22. Atamna H. and Ginsburg H. (1993) Origin of reactive oxygen species in erythrocytes infected with *Plasmodium falciparum*. Mol. Biochem. Parasitol. 61, 231–241

23. Vander Jagt D.L., Hunsaker L.A., Campos N.M., and Baack, B.R. (1990) D-lactate production in erythrocytes infected with *Plasmodium falciparum*. Mol. Biochem. Parasitol. 42, 277–284

24. Mehta, M., Sonawat, H. M., and Sharma, S. (2006) Glycolysis in *Plasmodium falciparum* results in modulation of host enzyme activities. J. Vector Borne Dis. 43, 95–103

25. Muralidharan, V., Oksman, A., Iwamoto, M., Wandless, T. J., and Goldberg, D. E. (2011) Asparagine repeat function in a *Plasmodium falciparum* protein assessed via a regulatable fluorescent affinity tag. Proc. Natl. Acad. Sci. 108, 4411–4416

26. Pei, Y., Miller, J. L., Lindner, S. E., Vaughan, A. M., Torii, M., and Kappe, S. H. (2013) Plasmodium yoelii inhibitor of cysteine proteases is exported to exomembrane structures and interacts with yoelipain-2 during asexual blood-stage development. Cell. Microbiol. 15, 1508–1526

27. Sievers, F., Wilm, A., Dineen, D., Gibson, T. J., Karplus, K., Li, W., Lopez, R., McWilliam, H., Remmert, M., Söding, J., et al. (2011) Fast, scalable generation of high-quality protein multiple sequence alignments using clustal omega. Mol. Sys. Biol. 7, 539

28. Smialowski, P., Doose, G., Torkler, P., Kaufmann, S., and Frishman, D. (2012) Proso ii-a new method for protein solubility prediction. FEBS J. 279, 2192–2200

29. Michaelis L. and Menten M. (1913) Die kinetik der invertinwirkung. Biochem Z 49, 333–369.

30. Godiska, R., Mead, D., Dhodda, V., Wu, C., Hochstein, R., Karsi, A., Usdin, K., Entezam, A., and Ravin, N. (2009) Linear plasmid vector for cloning of repetitive or unstable sequences in *Escherichia coli*. Nucleic Acids Res. 38, e88

31. Janse, C. J., Ramesar, J., and Waters, A. P. (2006) High-efficiency transfection and drug selection of genetically transformed blood stages of the rodent malaria parasite Plasmodium berghei. Nat. Protocols 1, 346

